# The *Freesia refracta* virome analysis sheds new light on the phylogenetic relationships in the *Konkoviridae* and *Yueviridae* families

**DOI:** 10.1101/2025.03.19.644121

**Authors:** Monica Marra, Silvia Rotunno, Fulco Frascati, Roberto Pierro, Pasquale Restuccia, John Hammond, Anna Maria Vaira, Laura Miozzi

## Abstract

The necrosis syndrome of freesia, first described in 1970 in Northern Europe, is still jeopardizing freesia cultivation all over the world. Although several viruses have been listed as possible causal agents, the etiology of the disease is still not clear and is possibly linked to a combination of different factors. In this study, a high-throughput sequencing virome analysis was performed on total RNA extracts derived from symptomatic freesia leaves; a novel virus putatively belonging to the recently ratified *Konkoviridae* family in the *Bunyaviricetes* class has been identified and characterized, for which we propose the name of freesia konkovirus 1 (FreKV-1). This family, officially listing only one genus and two species, has been expanded by exploring publicly available metatranscriptomic datasets through the Serratus Project Database and reconstructing new viral entities; the phylogenetic position of the *Konkoviridae* family has been investigated and new genera belonging to the family have been proposed. Moreover, a further previously unknown virus, putatively belonging to the *Yueviridae* family was partially characterized and its phylogenetic position was discussed. Overall, the analysis increased our knowledge of the number of viral agents infecting freesia and possibly involved in freesia necrosis syndrome.

## Introduction

The necrotic disorder of freesia (*Freesia refracta* hyb., family Iridaceae) was first described in the Netherlands more than 50 years ago and reported in Germany and England in the same period. By the beginning of this century this disease appeared to cause considerable economic losses in Italy and in the Netherlands. Typical symptoms are chlorotic inter-veinal spots and streaks at the leaf tips that expand downwards and turn necrotic and may vary according to cultivar and climate (Figure 1).

**Figure 1.**
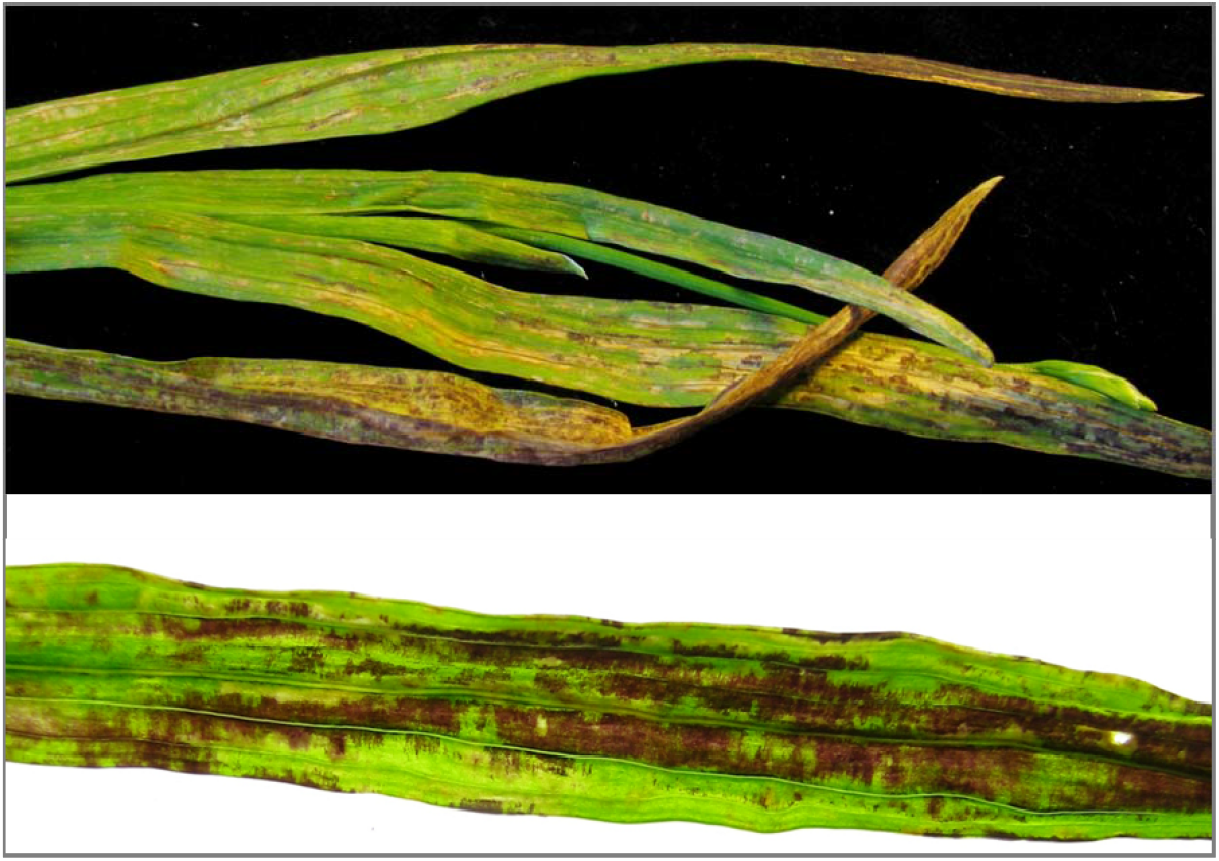
Necrotic and deformed freesia leaves showing typical symptoms of freesia necrotic disorder; details are shown on the lower panel.

In the past, the syndrome was supposed to be caused by a mixed virus infection that had among the main actors the potyvirus freesia mosaic virus (FreMV; *Potyvirus freesiae*) and a not yet well characterized varicosavirus (Vaira et al. 2015). Subsequently, the disease has been widely linked to the presence of a newly, at that time, discovered ophiovirus named freesia sneak virus (FreSV; *Ophiovirus freesiae*) (Rotunno et al. 2024), belonging to the family *Aspiviridae* (Kuhn et al. 2022). FreSV is soil□transmitted by *Olpidium* sp. (Meekes and Verbeek 2011) and has up to now been detected in freesia in several temperate world regions, including Europe, North America (Vaira et al. 2009), New Zealand (Pearson et al. 2009) and South Korea (Jeong et al. 2014). Until now FreSV has been reported only in freesia and in lachenalia (*Lachenalia* hyb., Hyacinthaceae) (Vaira et al. 2007) showing similar symptomatology. Even if the correlation between the presence of symptoms and FreSV detection is high, some uncertainty remains, and FreSV can be considered one of the main agents involved in the syndrome but not the only one.

Here, a high-throughput sequencing (HTS) analysis of total RNA extracted from symptomatic freesia leaves was performed in order to expand our knowledge on the virome associated with the necrotic disorder. Two new viral species representing new members in the *Konkoviridae* and in the *Yueviridae* families, respectively, were identified. Furthermore, the open-science viral discovery platform Serratus Project Database (Edgar et al. 2022) has been used to explore the virome associated to already available transcriptome sequencing data of millions of samples collected by the world biological community, and potential seven new members of the *Konkoviridae* family were discovered, thus expanding the newly created *Konkoviridae* family. Based on phylogenetic analysis, we propose here the reorganization of the *Konkoviridae* family adding two new genera and identifying a new *Konkoviridae* species isolated in freesia as representative of one of them.

## 2. Materials and Methods

### 2.1 Sample collection

Necrotic foliar tissue from freesia plants was collected in Liguria region (North-West Italy) in 2011 (sample fr231211) and 2014 (sample fr140314), both in Sanremo area in the same plots and used for HTS analysis. More recently, 22 plants showing various degrees of necrotic symptoms were collected in 2022 in the same area; 6 of them, particularly symptomatic, were pooled (sample fr090322) for HTS analysis. All samples were stored at −80°C until processing.

### 2.2 Sample processing and Illumina sequencing

Total RNA was individually extracted by Spectrum™ Plant Total RNA Kit (Sigma Aldrich, St. Louis, MO, USA) following manufacturer’s instructions. After DNase treatment with the Turbo DNA-free kit (Invitrogen, UK), total RNA extracted from samples collected from 2011 to 2014 was pooled and sent to the specialized company Macrogen (Seoul, Republic of Korea; http://www.macrogen.com) for Illumina TruSeq Stranded Total RNA Library construction with riboZero Plant ribosomal removal and HTS using the Illumina NovaSeq 6000 100bp paired-end sequencing system. RNA from samples collected in 2022 was pooled and sent to the specialized company Novogene (Cambridge, United Kingdom; https://www.novogene.com/), for HTS with Illumina NovaSeq 150bp paired-end sequencing system.

### 2.3 Virome annotation

Raw reads were processed with fastp v. 0.21.0 (Chen et al. 2018), using default parameters, for quality check and removal of adapters. Residual ribosomal sequences were discarded using bbduk script (default parameters) from bbmap v. 38.7 (Bushnell et al. 2017). Reads were then assembled using metaspades v. 3.15.1 (Nurk et al. 2017), using k-mer values of 71, 81 and 91, and obtained contigs were further extended with CAP3 (VersionDate: 02/10/15) (Huang and Madan 1999). Contigs with a length above 600 nt were queried against the NCBI non redundant protein database (https://www.ncbi.nlm.nih.gov/) using DIAMOND blastx with an e-value cutoff of 1e^-5^. In order to explore the taxonomical content, results from DIAMOND blastx were saved as data file and uploaded into MEGAN v. 6.24.1. Viral contigs identified using DIAMOND blastx were further queried against the NCBI non redundant nucleotide database (https://www.ncbi.nlm.nih.gov/) using blastn. ORFfinder (https://www.ncbi.nlm.nih.gov/orffinder/), with default parameters and a minimal ORF length of 300 nt was used to identify open reading frames and encoded protein sequences.

### 2.4 Molecular detection, RACE-PCR and validation

Diagnostic, validation and RACE primers used in this study are listed in Table S1. Virus-specific primers were synthesized matching the sequences obtained by HTS analysis. cDNA synthesis was performed using High-Capacity cDNA Reverse Transcription Kit (Thermo Fisher Scientific, Waltham, MA, USA) and PCR reactions were carried out using Platinum II Taq Hot-Start DNA Polymerase (Thermo Fisher Scientific, Waltham, MA, USA) according to manufacturers’ instructions. RT-PCR was performed in some cases using the faster procedure of the One-step RT-PCR kit (Qiagen, Hilden, Germany), following manufacturer’s instructions. PCR amplified fragments were visualized by electrophoresis using 1.5% agarose gel, purified with the Zymoclean DNA Clean & Concentrator Kit (Zymo Research, CA, USA) and sent to a specialized company (BMR Genomics, Padova, Italy) for Sanger sequencing. For RACE-PCR, the 5’/3’ RACE Kit (Roche, Basel, Switzerland) was used according to manufacturer’s instructions.

### 2.5 Metatranscriptomic data mining

Palmprints associated with putative novel konkoviruses were identified querying the Serratus Project Database using as input the RdRp protein sequence of freesia konkovirus 1. SRA raw data associated with selected palmprints were downloaded from the NCBI-SRA database, pre-processed using fastp v. 0.21.0 and *de novo* assembled using metaspades v. 3.15.1. Obtained contigs were compared using blastx with the protein sequences of the known konkoviruses (FreKV-1, TuSV, LBVaPV, LacPhV-1). ORFfinder (https://www.ncbi.nlm.nih.gov/orffinder/) was used to identify open reading frames and encoded protein sequences. Alignments were performed using MEGA11 v11.0.11.

### 2.6 Phylogeny reconstruction

Viral sequences were retrieved from the NCBI protein database (https://www.ncbi.nlm.nih.gov/). The updated taxonomy (Virus Taxonomy: 2023 Release) reported by the International Committee on Taxonomy of Viruses (ICTV Website, accessed on 10/24/2024) was used as a guide to retrieve sequences of each representative virus species. Sequences were aligned using MEGA11 (v11.0.11), applying the MUSCLE algorithm. Phylogenetic tree was generated with MEGA11 (v11.0.11), with the Maximum Likelihood (ML) method, applying the Jones-Taylor-Thornton (JTT) model.

## 3. Results

### 3.1 Overview of the virome associated with the necrotic disorder of freesia

Freesia leaves showing necrotic disorder symptoms were collected in Liguria (Northern Italy) in 2011 and 2014 and used for virome analysis through total RNA Illumina sequencing. After cleaning, a total of 47,516,574 clean reads were obtained and *de novo* assembled in 14,746 contigs (contig length >600 nt). Querying these contigs against the non-redundant protein NCBI database (https://www.ncbi.nlm.nih.gov/), a total of 8 contigs showing similarity (e-value < e^-5^) with viral protein sequences were identified (Table 1A and Supplementary Table S2). These contigs were further compared with nucleotide sequences available in the NCBI non-redundant nucleotide database. One contig (c1_11/14) showed a high level of identity with the polyprotein of the potyvirus FreMV. With a reads percentage of 2.36%, this contig was the most represented in terms of reads abundance. Four contigs (c2_11/14, c3_11/14, c4_11/14, c5_11_14) shared similarity with proteins encoded by the ophiovirus FreSV and corresponded to its four RNA genomic segments. In terms of read abundance, contigs c2_11/14, c3_11/14 and c4_11/14 were represented with a read percentage ranging from 0.29 to 0.15, while c5_11/14, corresponding to RNA4, was less represented (0.05% of total reads). Two contigs (c6_11/14, c7_11/14) showed a certain level of similarity with the RNA-dependent RNA polymerase and the nucleocapsid protein of lachenalia phenuivirus-1 (LacPhV-1), a newly identified viral species with similarity with tulip streak virus (TuSV), the first recognized member of the recently ratified *Konkoviridae* family (Dekker et al. 2024; Neriya et al. 2021; Neriya and Nishigawa 2024). In terms of read abundance, c6_11/14 was the most represented of the two (1.64%) while c7_11/14 was represented by 0.88% of total reads. Considering the level of similarity of our contigs with LacPhV-1, we proposed that they represent a distinct new viral species belonging to the *Konkoviridae* family and we will discuss it in detail in dedicated section 3.2. Finally, one contig (c8_11/14) shared a limited similarity at protein level with the putative RNA-dependent RNA polymerase of *Fusarium culmorum* yue-like virus 1 (FcYV1), belonging to the *Yueviridae* family. This contig had no significant similarity at the nucleotide level in the non-redundant nucleotide NCBI database and, in terms of reads abundance, was the least represented among all viral contigs, since only 0.003% of the total number of reads mapped on it, equally distributed in both plus and minus polarities. Contig c8_11/14 is likely to represent a new virus belonging to the *Yueviridae* family and will be discussed in dedicated section 3.4.

**Table 1.**
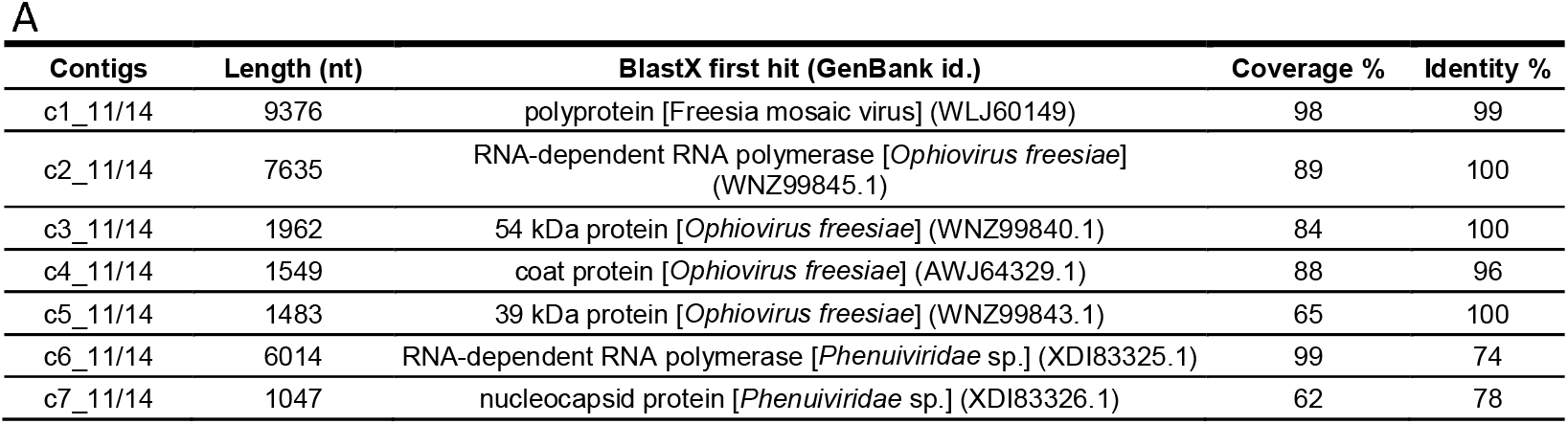
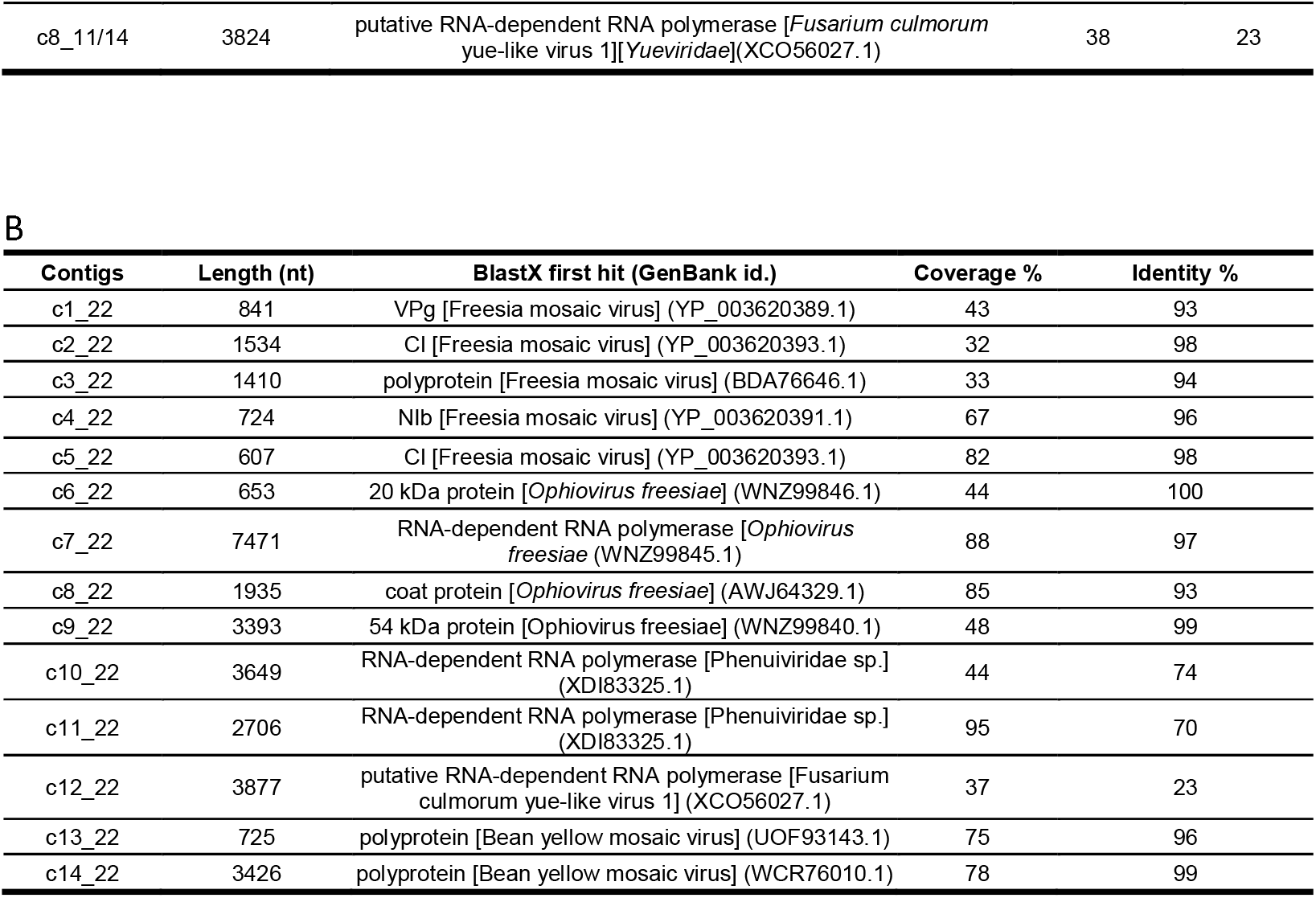
A) Viral contigs identified in freesia leaves showing necrotic disorder symptoms collected in Liguria (Northern Italy) during the 2011-2014 years; B) Viral contigs identified in freesia leaves showing necrotic disorder symptoms collected in Liguria (Northern Italy) during the 2022 year.

The presence of the viral entities identified in symptomatic freesia leaves was confirmed by an independent RNA-Seq analysis performed on a pool of symptomatic leaves collected from six symptomatic freesia plants in 2022, in the same area. In this new RNA-Seq analysis, we obtained a total of 101,460,426 clean reads that were assembled in a total of 79.864 contigs (length > 600 nt); among them, we retrieved contigs associated with all the viruses identified in the freesia symptomatic leaves collected in 2011-2014 (Table 1B and Supplementary Table S2); in detail, five contigs (from c1_22 to c5_22) were associated to FreMV, four contigs (from c6_22 to c9_22) corresponded to FreSV, two contigs (c10_22 and c11_22) showed similarity with LacPhV-1 and one contig (c12_22) had weak similarity with the putative RNA-dependent RNA polymerase of FcYV1 and corresponded to the new putative member of the *Yueviridae* family already identified in the previous RNA-seq analysis (see Supplementary Table S2 for details). Furthermore, in these samples, we also identified two contigs (c13_22 and c14_22) with high similarity with the polyprotein of the potyvirus bean yellow mosaic virus (BYMV), a virus that was not present in the samples collected in 2011 and 2014. The presence of viral RNAs in freesia symptomatic samples collected in 2022 was confirmed independently in each of the six plants pooled for the RNA-Seq analysis by RT-PCR with specific primers followed by Sanger sequencing of the amplified PCR fragments (Supplementary Figure S1).

Overall, the analysis of the virome associated with the freesia necrotic disorder led us to identify the presence of five different viral entities. Three of them, i.e. FreMV, FreSV and BYMV, consist of viruses already known to infect freesia (Sastry et al. 2019), while the remaining two represent new viruses belonging to the recently ratified *Konkoviridae* and *Yueviridae* families.

### 3.2 Characteristics of a putative novel *Konkovirus* species

Some hints about this further putative actor in the necrotic disorder of freesia stage, came out for the first time when following a Sequence Independent Amplification (SIA) procedure (Agindotan et al. 2010) on total RNA extracted from necrotic freesia leaves collected in Liguria (Northern Italy) during the 2011-2012 winter season. From this analysis, we obtained clones of unknown virus-like genome sequences (Vaira et al. 2018). This new virus was shown to be mechanically transmissible from symptomatic freesia leaves to the test plant *Nicotiana benthamiana* even if in this plant no symptoms were evident.

As reported in the previous section, RNA-Seq analysis allowed us to reconstruct two contigs (c6_11/14 and c7_11/14) corresponding to RNA1 and RNA2 of a putative new member of the *Konkoviridae* family (Order *Hareavirales*, Class *Bunyaviricetes*). Rapid Amplification of cDNA Ends (RACE-PCR) were performed to obtain the complete sequence of these genomic segments. The *Konkoviridae* family, ratified by the ICTV in 2023, lists, at the moment, a single genus (*Olpivirus*) represented by the negative-sense and soil-borne RNA virus TuSV originating from tulip (*Tulipa gesneriana* L.) and by the RNA virus lactuca big vein associated phlebovirus (LBVaPV, *Olpivirus lactucae*) isolated in lettuce (Schravesande et al. 2024) (ICTV website accessed on 03/05/2025). A putative third member provisionally named LacPhV-1 and showing similarity with TuSV and LBVaPV (Table 2A,B) has been recently identified in lachenalia (Dekker et al. 2024 bioRxiv preprint). Since the RNA genome of these viruses is composed of 4 RNAs, we decided to further search our data for contigs having similarity with these RNAs and we identified two more contigs corresponding to the full-length genomic segments RNA3 and RNA4.

**Table 2.**
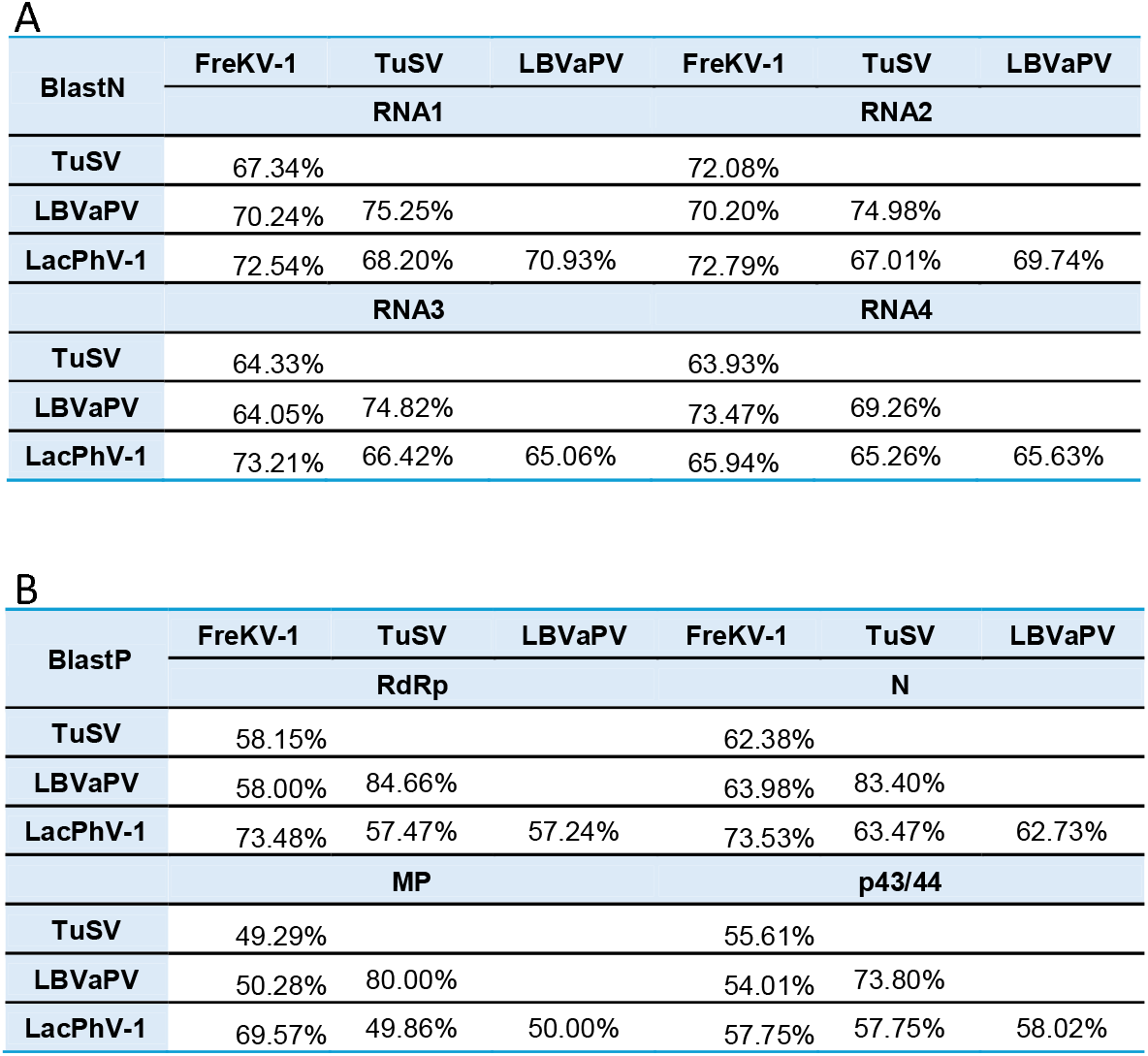
A) Nucleotide similarity among the RNA genome segments of FreKV-1 and the other two members of the konkoviridae family (TuSV, LBVaPV and LacPhV-1) either accepted or under evaluation; B) Amino acid similarity among the RNA genome segments of FreKV-1 and the other members of the konkoviridae family (TuSV, LBVaPV and LacPhV-1) either accepted or under evaluation; RdRp, RNA-directed RNA polymerase; N, nucleocapsid protein; MP, movement protein.

Overall, the complete genomic sequence of a novel konkovirus, constituted by four viral RNA segments (RNA1-RNA4; GenBank accessions: PQ490803, PQ490804, PQ490805, PQ490806) that code for four virus-related proteins (Figure 2 and Table 3) was reconstructed. These RNAs show similarity with the four genomic RNAs of TuSV, with identity percentage values ranging between 72% and 64% (Table 2) and coverage percentage ranging between 58% and 44% (Supplementary Table S3). We have identified four ORFs coding four proteins, i.e. the RNA-dependent RNA polymerase (RdRp), the nucleocapsid protein (N), the movement protein (MP) and the protein of unknown function p44 (Figure 2), that show an identity percentage ranging 62% and 58% and a coverage percentage ranging between 100% and 87% with the corresponding proteins encoded by TuSV (Table 2 and Supplementary Table S3). Since defective viral RNAs can have a role in modulating accumulation, infection dynamics and virulence of a virus (Budzyńska et al. 2022), it is worth noting that the RT-PCR with specific primers for the RNA4 (Supplementary Figure S1), followed by Sanger sequencing, highlighted the existence of a defective RNA4 of 829 nt with an ORF coding for a shorter version of the p44 protein. Further analysis would be useful to clarify the possible role of this defective RNA in the infection cycle.

**Table 3.** Full coding sequence organization of the four FreKV-1 genomic RNAs obtained in this study and statistics.

| Genomic segment | Length (nt) | n. of mapping reads | % of mapping reads | Average coverage | ORF sense | Protein function | Estimated molecular weight (kDa) / amino acids (aa) |
| --- | --- | --- | --- | --- | --- | --- | --- |
| RNA1 | 6339 | 792887 | 1.67 | 12479 X | vcRNA | RNA-dependent RNA polymerase | 239.2/2069 |
| RNA2 | 1042 | 417904 | 0.88 | 39964 X | vcRNA | Nucleocapsid protein | 27.7/239 |
| RNA3 | 1357 | 367182 | 0.77 | 27017 X | vcRNA | Movement protein | 40.0/354 |
| RNA4 | 1288 | 416758 | 0.88 | 32295 X | vcRNA | Unknown function | 44.2/374 |

**Figure 2.**
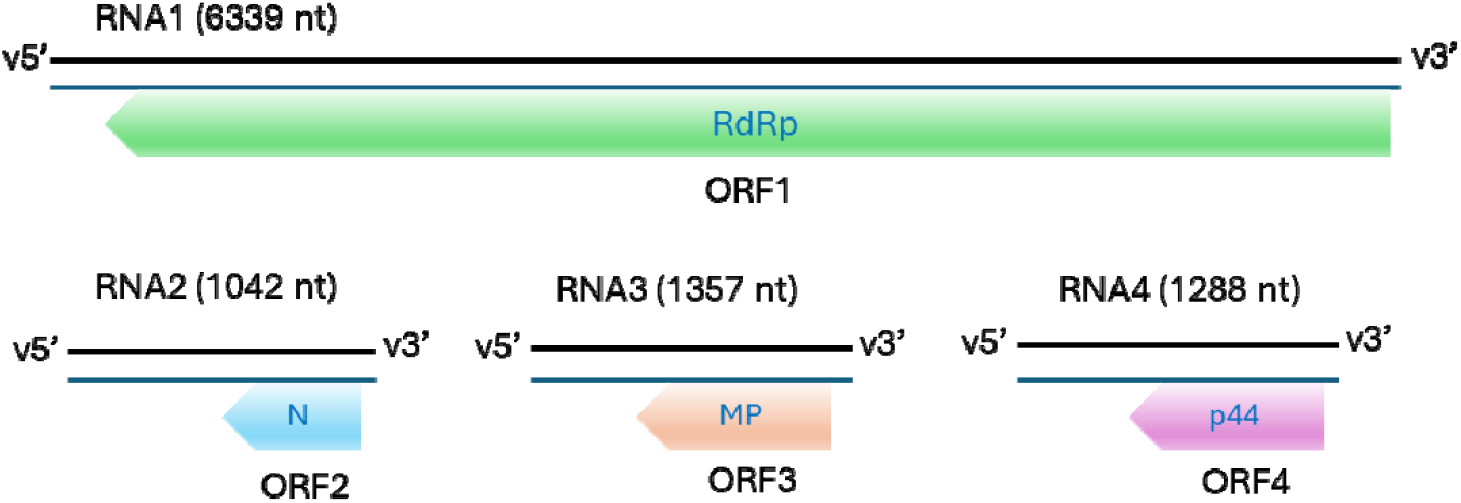
Genome organization of FreKV-1; RdRp, RNA-directed RNA polymerase; N, nucleocapsid protein; MP, movement protein

The identity percentage values at protein and nucleotide levels observed between our new virus, TuSV, LBVaPV and the other putative konkovirus LacPhV-1 (Table 3), and, in particular, the identity percentages of the RdRp protein sequences, support its classification as a distinct new species belonging to the new *Konkoviridae* family. Since the new virus was identified in freesia plants, we propose the name of Freesia konkovirus 1 (FreKV-1). Interestingly, these four viruses were found in plants from four different families: *Iridaceae* (FreKV-1), *Hyacinthaceae* (LacPhV-1), *Liliaceae* (TuSV) and *Asteraceae* (LBVaPV), suggesting that the host range of the konkoviruses is quite wide.

The sequences of both 5’ and 3’ ends of FreKV-1 genomic RNAs show characteristics typical for *Bunyaviricetes* (Reguera et al. 2013; Sasaya et al. 2023; Takahashi et al. 1990). In detail, like other members of the *Konkoviridae* family, the 5’ and 3’ terminal regions can form a panhandle structure (Figure 3A) and have almost identical sequences, with 11 and 13 conserved nucleotides, respectively (Figure 3B). The only exception is RNA4 that shows a reduced level of conservation at the 3’ termini.

**Figure 3.**
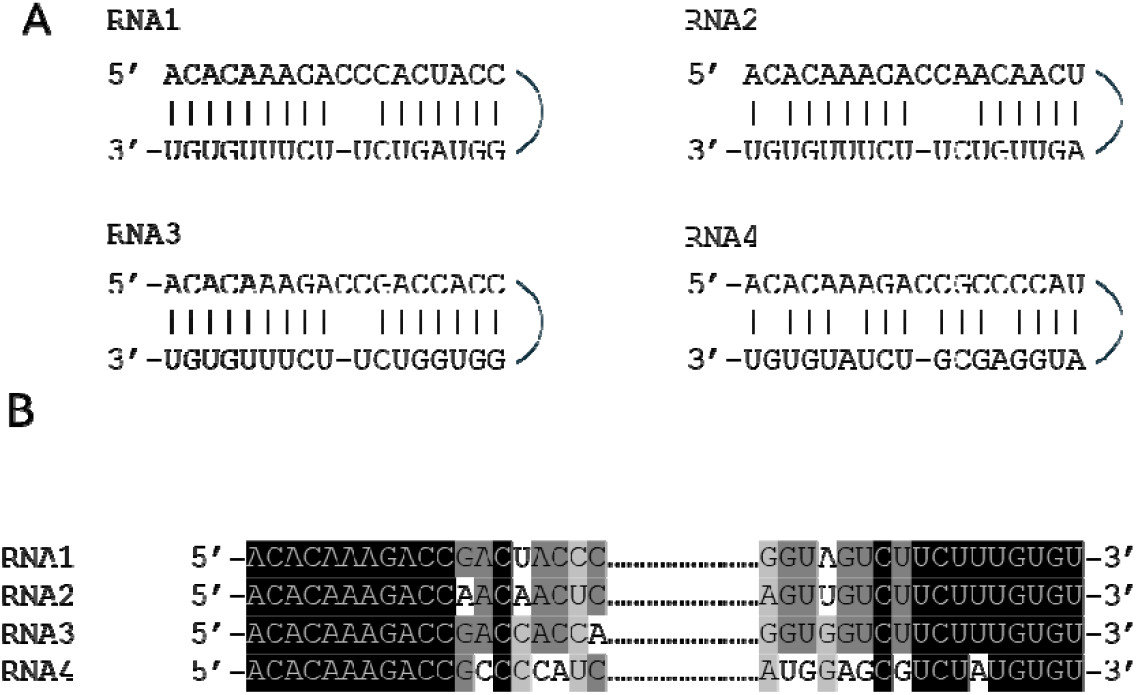
Analyses of FreKV-1 genomic RNA terminals. A) Alignment of FreKV-1 terminal sequences of genomic RNAs. B) Panhandle structures formed by the 5’ and 3’ termini of FreKV-1 genomic RNAs. Black, dark gray, light gray background corresponds to bases consensus to 4, 3, and 2 RNA segments, respectively

Comparing the terminal sequences of the four RNAs of FreKV-1 with those of the other konkoviruses, we observed that the RNA1 exhibits the higher level of conservation; indeed, the first 18 nt at 5’ termini and the first 17 nt at the 3’ termini of RNA1 were conserved between FreKV-1 and LacPhV-1 and only one nucleotide at each termini differentiated these viruses from LBVaPV and TuSV that, between them, share 100% identity at their RNA1 termini. All the other 5’ and 3’ termini were conserved for the first 11 to 14 nt with the only exception of the RNA4 that showed a relatively lower level of conservation, particularly in the case of FreKV-1 (Figure 4).

**Figure 4.**
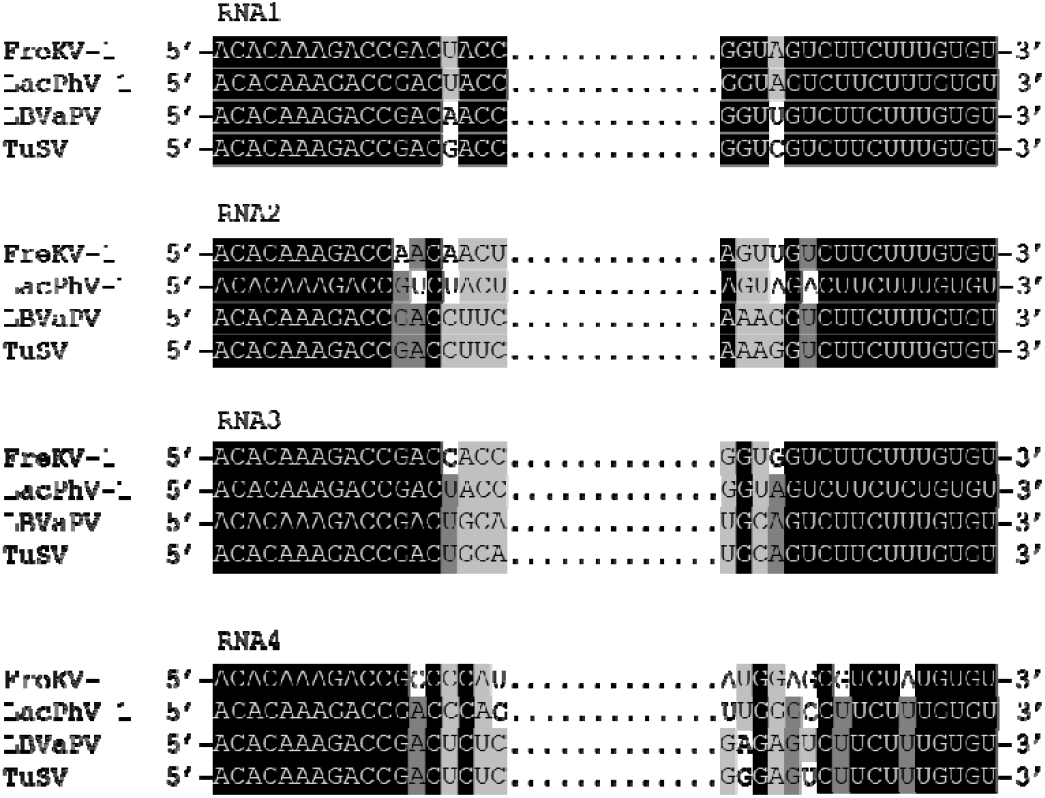
Alignment of genomic RNA terminals of FreKV-1, LacPhV-1, LBVaPV and TuSV, all official/putative members of the *Konkoviridae* family. Black, dark gray, light gray background corresponds to bases consensus to 4, 3, and 2 RNA segments, respectively

### 3.3 Metatranscriptomic exploration and phylogeny of the *Konkoviridae* family

Public databases contain a planetary collection of metatranscriptomic data that can be useful to reconstruct new viral genomes and extend our knowledge of the viral families. Here, we took advantage of the cloud computing infrastructure Serratus (Edgar et al. 2022) to search publicly available SRA-NCBI datasets (https://www.ncbi.nlm.nih.gov/sra/) for the presence of RdRp amino acid sub-sequences called ‘palmprint’ delineated by three essential motifs that together form the catalytic core in the RdRP structure. Querying the RdRp palmprint Serratus Database with the RdRp amino acid sequence of FreKV-1, we identified nine palmprints with an identity percentage greater than 70% (Figure 5 and Table 4). Among them, the palmprint u28439 corresponds to the RdRp of TuSV while the remaining eight are representative of new putative RNA viruses and, given the level of similarity, are representative of putative new members of the *Konkoviridae* family.

**Table 4.**
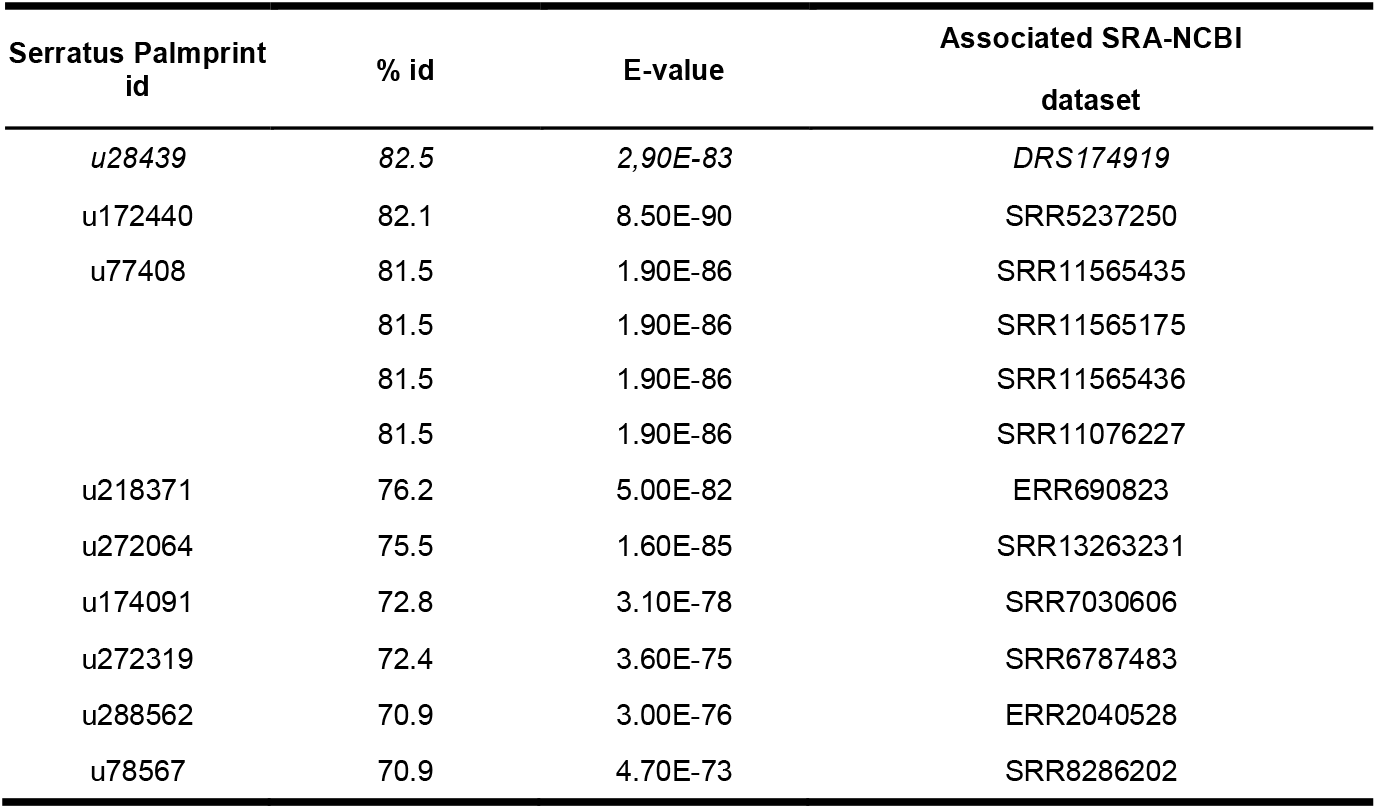
Serratus palmprints showing an identity percentage value above 70% with the RdRp FreKV-1 palmprint aminoacidic sequence; the palmprint corresponding to TuSV is in italic.

**Figure 5.**
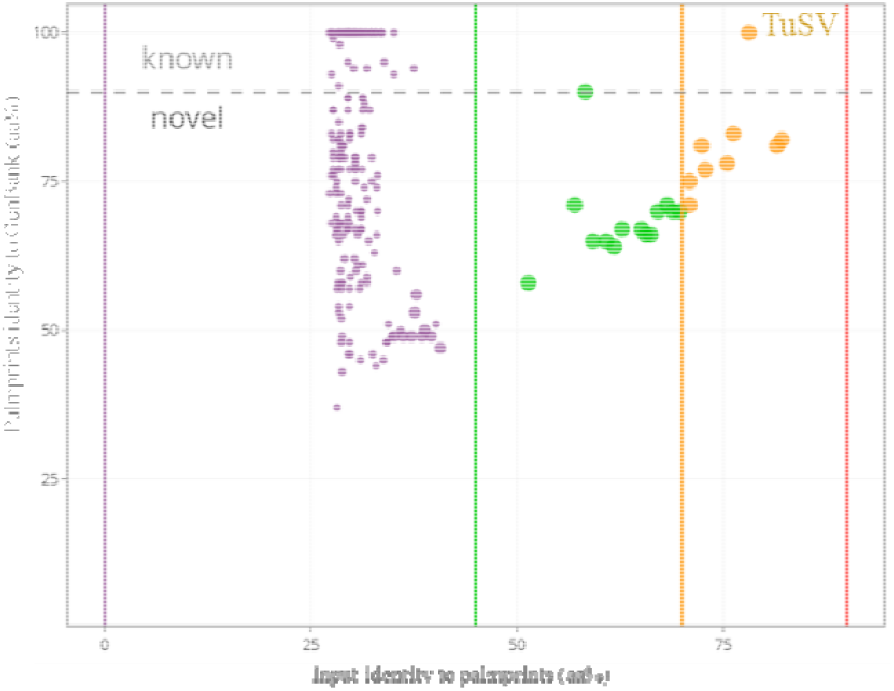
Percentage of identity of the RdRp FreKV-1 palmprint aminoacidic sequence used as input sequence with the palmprints in the Serratus Database. Green, orange and red lines indicate the 45%, 70% and 95% identity threshold, respectively. Dotted line corresponds to the 90% identity threshold and separates palmprints associated to known viruses (above the dotted line) and palmprints representing putative new viruses (below the dotted line). The figure was obtained by the Serratus Project Database tool.

To reconstruct the genome of the new konkoviruses identified by the palmprint similarity analysis, the raw reads of the selected SRA-NCBI datasets (Table 4) were retrieved, cleaned and *de novo* assembled, following the same bioinformatic pipeline used for the previous analyses; obtained contigs were then compared with the protein sequences of the four known konkoviruses (FreKV-1, TuSV, LBVaPV, LacPhV-1) using blastx; contigs representing viral RNA genomic sequences of new putative konkoviruses were identified and checked for their completeness. For seven out of the eight new putative konkoviruses, this approach allowed us to reconstruct the complete (or almost complete) CDS of RNA1, coding for the RdRp protein and RNA2, coding for the N protein; for two of them we were able to reconstruct the complete CDS of all the four expected RNA genomic segments (Table 5). It is worth noting that in the SRA-NCBI dataset SRR7030606, two different putative RNA1 complete CDSs were retrieved (blastx comparison: 53.27% identity-96% coverage and 43.95% identity-93% coverage in respect to the RdRp of FreKV-1), suggesting the existence of two different konkoviruses in this sample. In line with the hypothesis that these sequences correspond to new members of the *Konkoviridae* family and therefore are expected to infect plants, seven of them originated from SRA-NCBI datasets obtained from RNA extracted from plants belonging to different families that can be considered as putative hosts: *Waitzia nitida* (*Asteraceae*), *Campanula rotundifolia* (*Campanulaceae*), *Lilium leucanthum* (*Liliaceae*), *Epipactis purpurata* (*Orchideaceae*), *Tripterocalyx crux-maltae* (*Nyctaginaceae*), *Lennoa madreporoides* (*Lennoaceae*), *Erythranthe pardalis* (*Phrymaceae*), *Holcus lanatus* (*Poaceae*) (Table 5). The remaining palmprint originated from SRA-NCBI datasets is associated with soil metatranscriptomic data, in line with the supposed soil-borne nature and fungal mediated transmission of the viral members of the *Konkoviridae* family (Neriya et al. 2021). Based on the plants from which the SRA-NCBI datasets originated, for the new konkoviruses we proposed the following names: Waitzia associated konkovirus 1 (WaKV-1), Campanula associated konkovirus 1 (CaKV-1),Lilium associated konkovirus 1 (LilaKV-1), Epipactis associated konkovirus 1 (EpaKV-1), Epipactis associated konkovirus 2 (EpaKV-2), Tripterocalyx associated konkovirus 1 (TaKV-1), Lennoa associated konkovirus 1 (LeaKV-1), Erythranthe associated konkovirus 1 (EryaKV-1). For the new konkovirus identified in the metatranscriptomic soil we propose the provisional name of soil associated konkovirus (SaKV).

**Table 5.** Novel putative konkoviruses identified by bioinformatic analysis. * indicates genomic sequences with partial CDS. ^§^ Putative host was inferred from NCBI Biosample origin

| Novel konkovirus | Serratus Palmprint id | Associated SRA-NCBI dataset | Associated NCBI Biosample | Putative host<sup>s</sup> | Genomic segments recovered (length) |
| --- | --- | --- | --- | --- | --- |
| *Waitzia* associated konkovirus 1 (WaKV-1) | u172440 | SRR5237250 | SAMN06309684 | *Waitzia nitida* | RNA1 (6381 nt)<br>RNA2 (1324 nt)<br>RNA3 (1587 nt)<br>RNA4 (1307nt) |
| Soil associated konkovirus (SaKV) | u77408 | SRR11565175 | SAMN14514881 | --- | RNA1 (6356 nt)<br>RNA2 (1152 nt)<br>RNA3 (1003 nt)<br>RNA4 (1342 nt) |
| *Campanula* associated konkovirus 1 (CaKV-1) | u218371 | ERR690823 | SAMEA3157961 | *Campanula* aff. *rotundifolia* | RNA1* (1514 nt)<br>RNA2 (1010 nt)<br>RNA3 (1294 nt) |
| *Lilium* associated konkovirus 1 (LilaKV-1) | u272064 | SRR13263231 | SAMN17082061 | *Lilium leucanthum* var. *centifolium* | RNA1* (875-752-617-470 nt),<br>RNA2* (295 nt)<br>RNA3* (438-279 nt)<br>RNA4* (222 nt) |
| *Epipactis* associated konkovirus 1 (EpaKV-1) /<br>*Epipactis* associated konkovirus 2 (EpaKV-2) | u174091 | SRR7030606 | SAMN08949893 | *Epipactis purpurata* | RNA1 (6508 nt)<br>RNA1 v2 (6306 nt)<br>RNA2 (1165 nt) |
| *Tripterocalyx* associated konkovirus 1 (TaKV-1) | u272319 | SRR6787483 | SAMN07527256 | *Tripterocalyx crux-maltae* | RNA1 (6322 nt)<br>RNA2 (1038 nt)<br>RNA3 (1375 nt) |
| *Lennoa* associated konkovirus 1 (LeaKV-1) | u288562 | ERR2040528 | SAMEA104170572 | *Lennoa madreporoides* | RNA1 (5459 nt)<br>RNA2 (778 nt) |
| *Erythranthe* associated konkovirus 1 (EryaKV-1) | u78567 | SRR8286202 | SAMN10527072 | *Erythranthe pardalis* | RNA1 (5087 nt)<br>RNA2 (1053 nt) |

The representatives of all thirty-seven genera of viruses included in the eight families of the *Hareavirales* order were used to infer the phylogenetic relationships of the RdRp and N proteins obtained from the already known and newly identified putative konkoviruses. The maximum likelihood (ML) tree of the RdRp proteins (Fig. 5) showed that all the konkovirus RdRp sequences group together in a monophyletic group, clearly separated from the other families of the *Hareavirales* order. The existence of this group is also highlighted by the identity percentage of the RdRp sequences as highlighted in the similarity matrix (Supplementary Figure S2). The same group is also evident in the phylogenetic tree obtained with the N protein sequences and in the corresponding similarity matrix (Supplementary Figure S3 and S4). The phylogenetic analysis of the RdRp protein sequences suggests the existence of at least three distinct genera in the *Konkoviridae* family: the official genus *Olpivirus*, represented by TuSV (*Olpivirus tulipae*) and LBVaPV (*Olpivirus lactucae*), a new one represented by the newly identified FreKV-1 (this study), LacPhV-1(Dekker et al. 2024), WaKV-1 (this study) and SaKV (this study) and a third one including LeaKV-1, EryaKV-1 and EpaKV-1 (Fig. 5).

**Figure 5.**
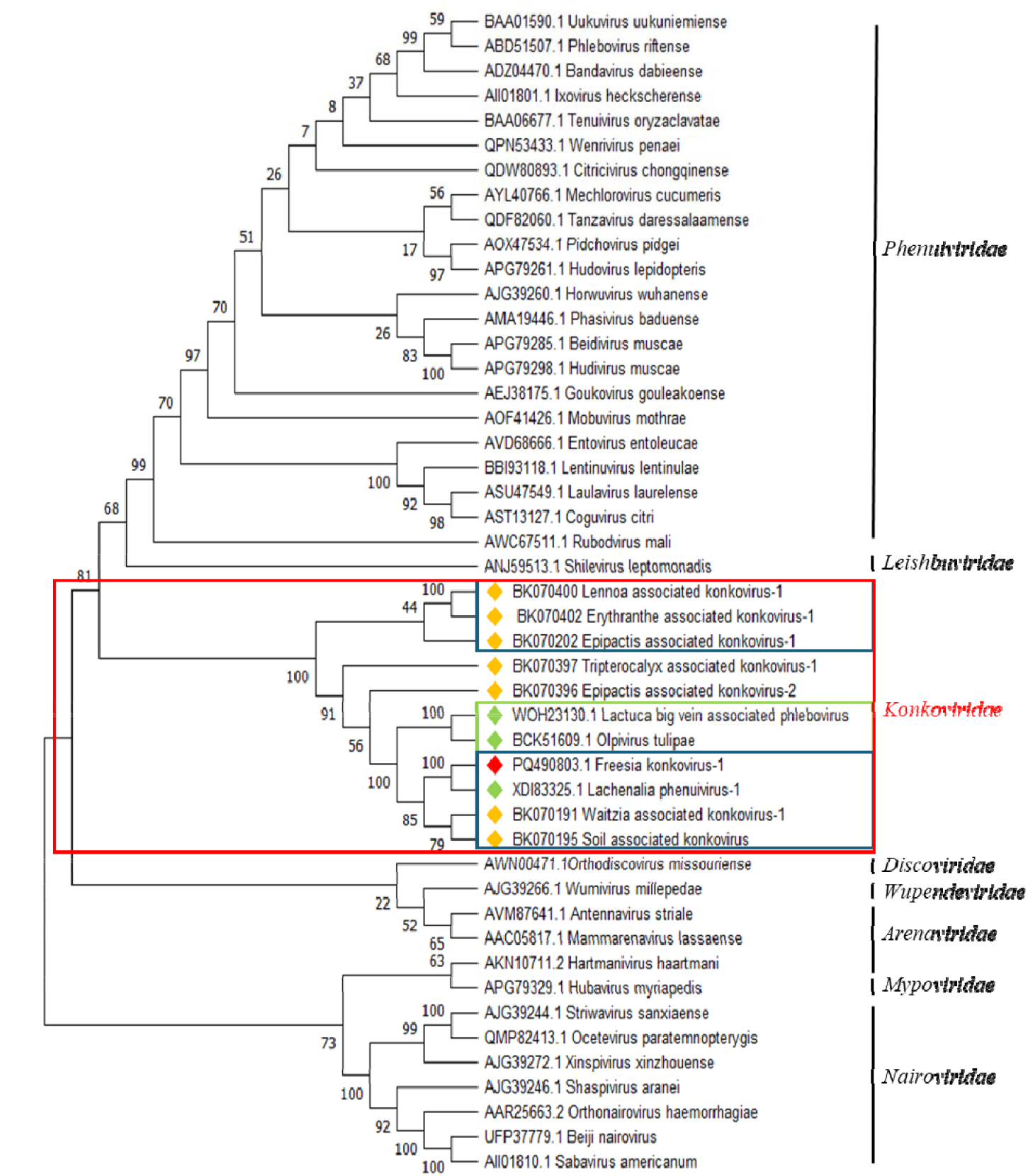
Phylogenetic tree of replicase aminoacid sequences of selected viral species listed in the *Hareavirales* order, the official species in *Konkoviridae* family and the newly discovered species possibly listing in *Konkoviridae* family (for which the RNA-1 full CDS has been obtained), computed by maximum likelihood (JTT matrix-based method). Consensus tree is constructed from 1,000 bootstrap trees. Numbers at nodes represent the percentage bootstrap values. Red diamond indicates FreKV-1, green diamonds indicate the already known konkoviruses including LacPhV-1 not yet ratified as member of the *Konkoviridae* family, orange diamonds indicate putative new members of the *Konkoviridae* family identified through the Serratus Project Database and reconstructed by bioinformatic analysis. Blue rectangles indicate the proposed new genera in the *Konkoviridae* family; the green rectangle indicates the only official genus ratified in the *Konkoviridae* family according to ICTV

### 3.4 Characteristics and phylogeny of a putative novel *Yueviridae* species

Analyzing the RNA-seq data, we also retrieved a contig of 3824 nt having highest identity to the putative RNA-dependent RNA polymerase of *Fusarium culmorum* yue-like virus 1, *Yueviridae* family (GenBank id. XCO56027.1) as first BLASTX hit, with 22.71% of protein identity and 38% coverage. The integration of this viral sequence in the genomic freesia DNA was excluded by PCR versus RT-PCR test (data not shown). Furthermore, we verified the existence of HTS reads mapping on both positive and negative senses of the viral genomic sequence as indication of possible dsRNA accumulation, typical of viral replication. Based on these data, we concluded that this contig represents a replicating RNA segment corresponding to a new yue-like virus for which we proposed the provisional name of freesia associated yue-like virus 1 (FraYV-1). *Yueviridae* (https://ictv.global/report/chapter/yueviridae/yueviridae) is a family of negative-sense RNA viruses belonging to the order *Goujianvirales* (*Yunchangviricetes* class, *Haploviricotina* subphylum); viruses of this family have bi-segmented negative single-stranded RNA genomes, with each segment possessing at least one ORF. The ORF in the L segment encodes a large protein corresponding to a RdRP and containing a GDP:polyribonucleotidyltransferase (PRNTase) domain related to that of viruses in the order *Mononegavirales*, suggesting a similar capping mechanism. The ORF in the S segment encodes a nucleocapsid protein structurally related to the homologs encoded by other members of the *Mononegavirales*, in particular paramyxovirids. The Family has been very recently established, and several biological aspects remain to be elucidated. At the moment, it includes a single genus with only two species, i.e. Běihăi sesarmid crab virus 3 (BhSCV3; *Yuyuevirus beihaiense*) and Shāhé yuèvirus-like virus 1 (ShYLV1; *Yuyuevirus shaheense*), which have been found in unspecified sesarmid crabs and freshwater isopods, respectively (Shi et al. 2016).

However, several unclassified yue-like viruses, often represented only by the L segment, have been also reported, associated with different hosts such as crustaceans and insects (Chiapello et al. 2021), stramenopiles (Charon et al. 2021; Chase et al. 2021; Chiapello et al. 2020), tracheophytes (Mifsud et al. 2023), and unspecified macrophytes (Rosario et al. 2022), thus suggesting that the diversity of *Yueviridae* is likely underestimated (Krupovic et al. 2023; Käfer et al. 2019; Olendraite et al. 2023; ICTV Website accessed on 10/23/2024). FraYV-1 is the only yue-like virus infecting angiosperms reported until now, thereby expanding the already vast host range of this family. The phylogenetic analysis of FraYV-1 highlighted the existence of two separate clades in the *Yueviridae* family; clade A includes the two yueviruses species defined in the ICTV taxonomy while clade B includes FraYV-1 and yue-like viruses isolated from different oomycetes and from fern (Fig. 6A). The catalytic domain in the conserved motif C of the RdRp proteins (Venkataraman et al. 2018) belonging to clade A shows the canonical GDD sequence while all the RdRp proteins belonging to clade B houses the IDD sequence instead. In this motif, a conserved G is present after DD in all the yueviridae members (Fig. 6B). These data suggest that the *Yueviridae* family could be split into two different genera and that further investigations are needed to better understand the phylogenetic relationships of the members of this family.

**Figure 6.**
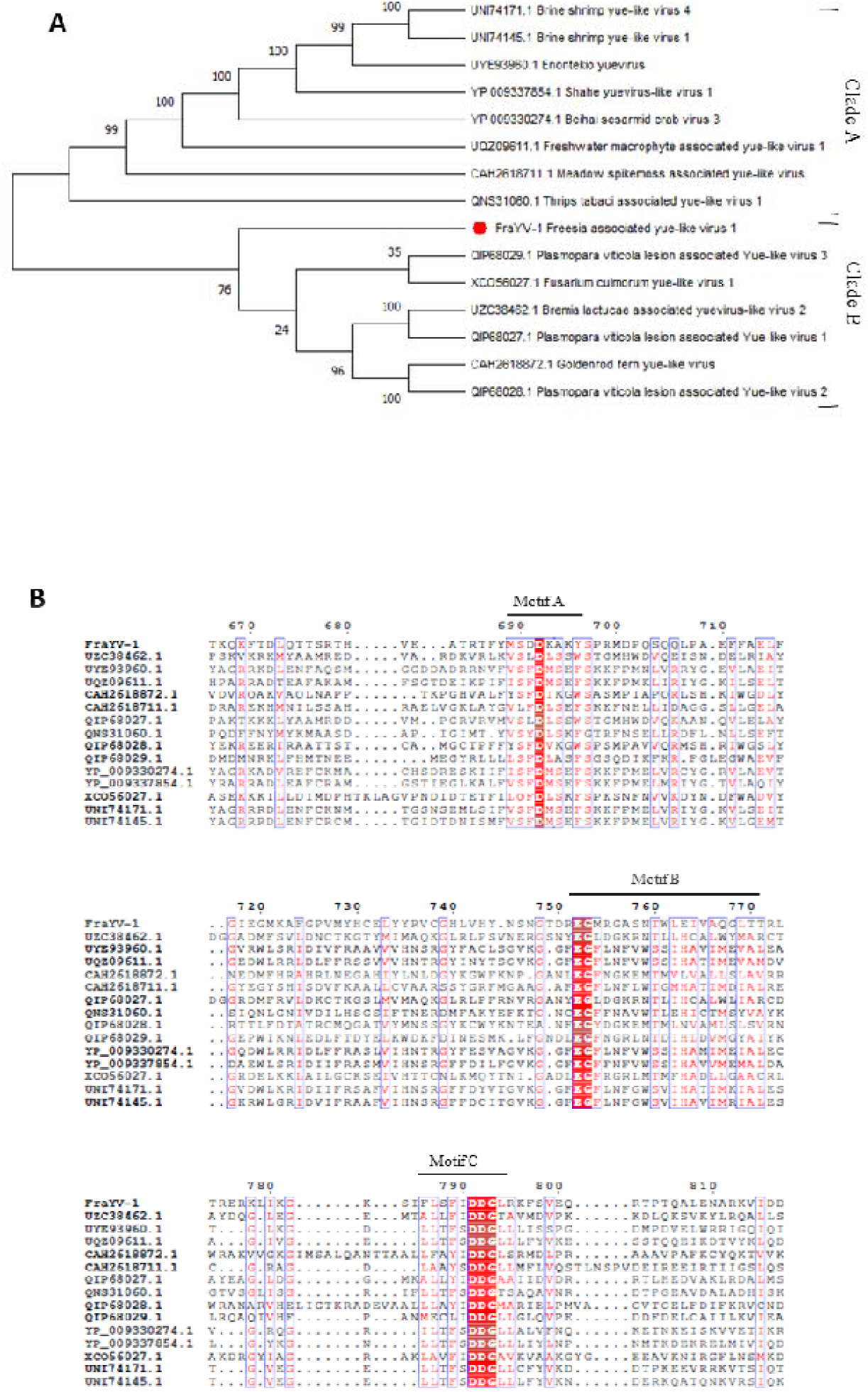
A) Phylogenetic tree of the *Yueviridae* family computed by maximum likelihood (JTT matrix-based method). Consensus tree is constructed from 1,000 bootstrap trees. Number at nodes are the percentage bootstrap values. Red diamond indicates FraYV-1; B) Alignment of RdRp protein sequences of the members of the *Yueviridae* family.

### 3.5 Virus detection and symptomatology

In order to clarify the possible correlation among the identified viruses and the necrotic disorder of freesia, we further investigated the level of association between the identified viruses and the necrotic symptomatology. Twenty-one freesia plant samples (1-21) showing different types of necrosis and an asymptomatic plant used as negative control (22), collected in 2022 in Northern Italy, have been tested individually using specific primers to detect the viruses revealed from the virome analysis (Figure 7). The potyvirus FreMV, not associated with necrosis in literature (Sastry et al. 2019), was present in all samples, including samples 1 and 9 in which some necrosis was only slightly evident on the collected leaves, and in the asymptomatic sample 22. BYMV was detected in almost all samples including the asymptomatic sample 22, with the only exception of samples 2, 9 and 19. Although this virus was widely reported in freesia, it has been mainly associated with chlorotic leaves, corm necrosis and color breaks in flowers, but was never clearly associated with the necrotic disorder. Our results indicate that, although both FreMV and BYMV may be widely spread in freesia cultivations, they are not unequivocally correlated to the presence of necrotic symptoms and therefore confirm that neither FreMV nor BYMV can be considered the etiological agent of the necrotic disorder of freesia. The ophiovirus FreSV, previously proposed as the causal agent but never convincingly associated with the disease, was detected in 15 out of 22 samples and the newly identified FreKV-1 in 6 out of 22; they were not diagnosed in the necrosis-free samples, but nevertheless the results suggest that neither virus appears to be itself the causative agent of necrosis. The presence of a wavy appearance of the foliar edge is prominent in plants positive for FreKV-1 and might be a symptomatology linked to its infection; nevertheless, a wider statistical analysis and biological assays are needed to confirm this hypothesis. The newly discovered FraYV-1 was found in 15 out of 22 samples, including the little symptomatic samples Fr1 and Fr9 and the asymptomatic one Fr22, revealing a high percentage of infection in freesia but not any clear association with the necrotic symptomatology.

**Figure 7.**
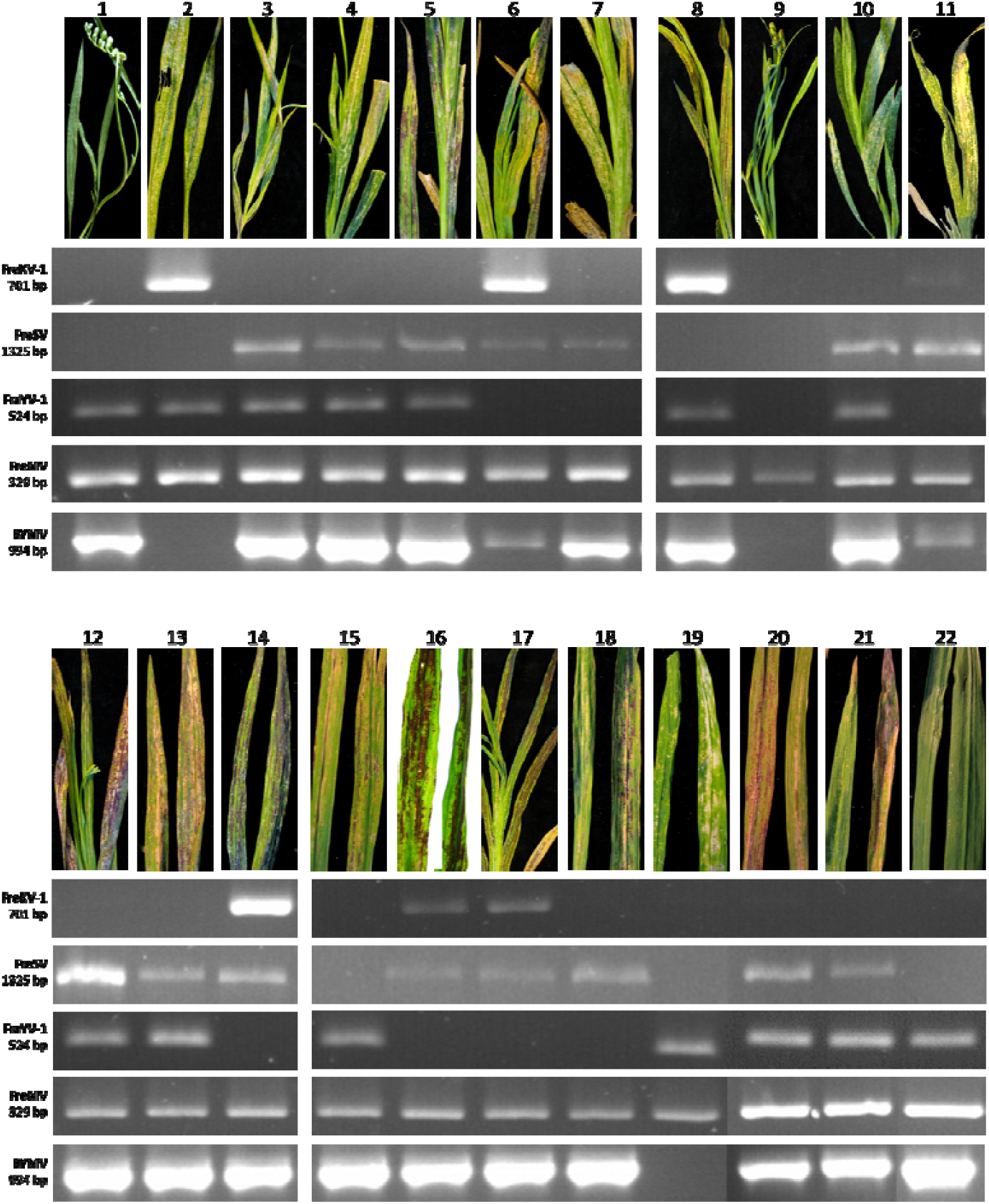
PCR detection of FreKV-1, FreSV, FreYV-1, FreMV and BYMV in single freesia plants showing various degrees of necrotic disorder symptoms

In the recent review about viruses infecting freesia, (Sastry et al. 2019) seven different viral species are listed. Three of them have been also found in the present investigation, the two potyviruses BYMV, FreMV and the ophiovirus FreSV, all considered widespread in freesia cultivations. In addition, the orthotospovirus Impatiens necrotic spot virus (INSV) has been reported, producing symptoms of chlorotic and necrotic leaf streaking in freesia; but it was not found in necrotic freesia leaves collected in this study. The same for the cucumovirus cucumber mosaic virus (CMV) and the tobravirus tobacco rattle virus (TRV), also not found in this study, that produce stunting of plants, chlorosis of leaves and deformation of flowers and are not directly linked with freesia necrotic disorder. Another viral agent, the varicosavirus freesia leaf necrosis virus (FrLNV), the somewhat elusive soil-born viral agent, previously most linked to the necrotic disorder of freesia, is again not found in necrotic freesia plants analyzed for this study. This complex scenario does not allow to clearly identify a single virus as the etiological agent of the severe leaf necrosis disorder, as it was called in the early studies, suggesting, once again, that the disease could arise from a combination of factors.

## 4. Discussion

In this study, the virome associated with symptomatic freesia plants was investigated. The plants exhibited varying degrees of necrosis, a disorder whose pathogenic agent remains largely unidentified. In the effort of identifying the etiological agent of the disorder, by using high-throughput sequencing technologies, we identified viral sequences corresponding to two previously uncharacterized viral entities belonging to the *Konkoviridae* and *Yueviridae* families, both of which were recently ratified by the ICTV. These families are known to contain only a small number of members, and data on them are very limited.

Our work took advantage from the use of high-throughput sequencing technologies that have revolutionized the study of viral biodiversity, greatly enhancing our understanding of viral ecosystems. Indeed, the reduction in sequencing costs, coupled with the advancement of high-performance sequencing technologies, has made these tools indispensable, which will continue to shape future virological research. However, it is worth underlining that these technologies have already generated a vast amount of data that are publicly available and that can be mined for valuable insights and that the exploration of these publicly available metatranscriptomic data offer enormous chances to explore the Earth RNA virome (Neri et al. 2022).

Here, we leveraged data of the Serratus Project Database (Edgar et al. 2022), specifically related to RdRp palmprints of RNA viruses, to identify potential new members of the *Konkoviridae* family and thereby expand our understanding of this family, very recently reported and listing only one official genus with two species. This approach facilitated the reconstruction of genomes (partial or complete coding sequences) for seven novel konkoviruses and the phylogenetic analysis of these sequences suggested the presence of at least three distinct genera within the family. The analysis of public datasets also broadened the host range of konkoviruses, suggesting their presence in a variety of plant species belonging to different plant families, both monocots and dicots, among which valuable ornamental plants are listed.

The HTS approach also enabled the identification of a novel virus with significant sequence similarity to members of the *Yueviridae* family. Phylogenetic analysis and conserved motif examination of the RdRps suggested that this family, which currently includes only two recognized members, could be split into two distinct clades, each potentially representing separate genera. Given the limited level of similarity among RdRp protein sequences, further investigations may even suggest the reclassification of these clades into two separate families: one comprising the two already classified yueviruses, and the other including yue-like viruses that infect oomycetes and plants, such as FraYV-1. The identification of additional yue-like viral sequences together with a better understanding of the biology of these viruses could help in deciphering their phylogenetic relationships. Furthermore, this is the first report of a yue-like virus infecting the clade of Angiospermae thus expanding the host range of this group of viruses.

The application of HTS once again underscored the complexity and underexplored nature of viral biodiversity. While this approach has been proved to be highly effective, it did not lead to definitively identifying the etiological agent of the necrotic disorder in freesia, likely because the symptoms are not solely attributable to a single pathogenic agent but being the result of the interplay of various biotic and abiotic factors that remain to be fully elucidated.

## 5. Conclusions

In summary, this work started as a study dedicated to the complex characterization of the virome associated with the freesia affected by the necrotic disorder ended in an interesting analysis of unknown viromes. Through a combined wet-lab and *in silico* approach, we improved the current knowledge of the diversity of viruses infecting freesia and contributed to expanding and clarifying the phylogenesis of the *Konkoviridae* and *Yueviridae* families, two clades ratified very recently and whose biodiversity is still largely underestimated.

## Supplementary data

Supplementary data are available at Virus Evolution online.

## Code and data availability

Raw reads of HTS related to this work are deposited in the Sequence Read Archive (SRA) of NCBI (https://www.ncbi.nlm.nih.gov/sra) with BioProject ID PRJNA1014396 and BioProject ID PRJNA1189045. Genomic sequence of known and new viruses identified in this study are deposited at NCBI with the following accession numbers: PV159347, PV171508, PQ490803, PQ490804, PQ490805, PQ490806, BK070191, BK070192, BK070193, BK070194, BK070195, BK070196, BK070197, BK070198, BK070199, BK070200, BK070201, BK070202, BK070203, BK070396, BK070397, BK070398, BK070399, BK070400, BK070401, BK070402, BK070403.

## Acknowledgments

This work is supported by EVA-Global project, funded by the European Union’s Horizon 2020 research and innovation program under grant agreement No. 871029 and by the European Commission— NextGenerationEU, Project SUS-MIRRI.IT “Strengthening the MIRRI Italian Research Infrastructure for Sustainable Bioscience and Bioeconomy”, code n. IR0000005. We would also thank Zocca Elena, Marian Daniele and Bordone Luca for precious greenhouse and laboratory work. Furthermore, we are grateful for the samples of diseased freesia provided by farms located in Sanremo region (Italy).

