## Supplementary Figure S1 for "The *Freesia refracta* virome analysis sheds new light on the phylogenetic relationships in the *Konkoviridae* and *Yueviridae* families"

### Slide 1
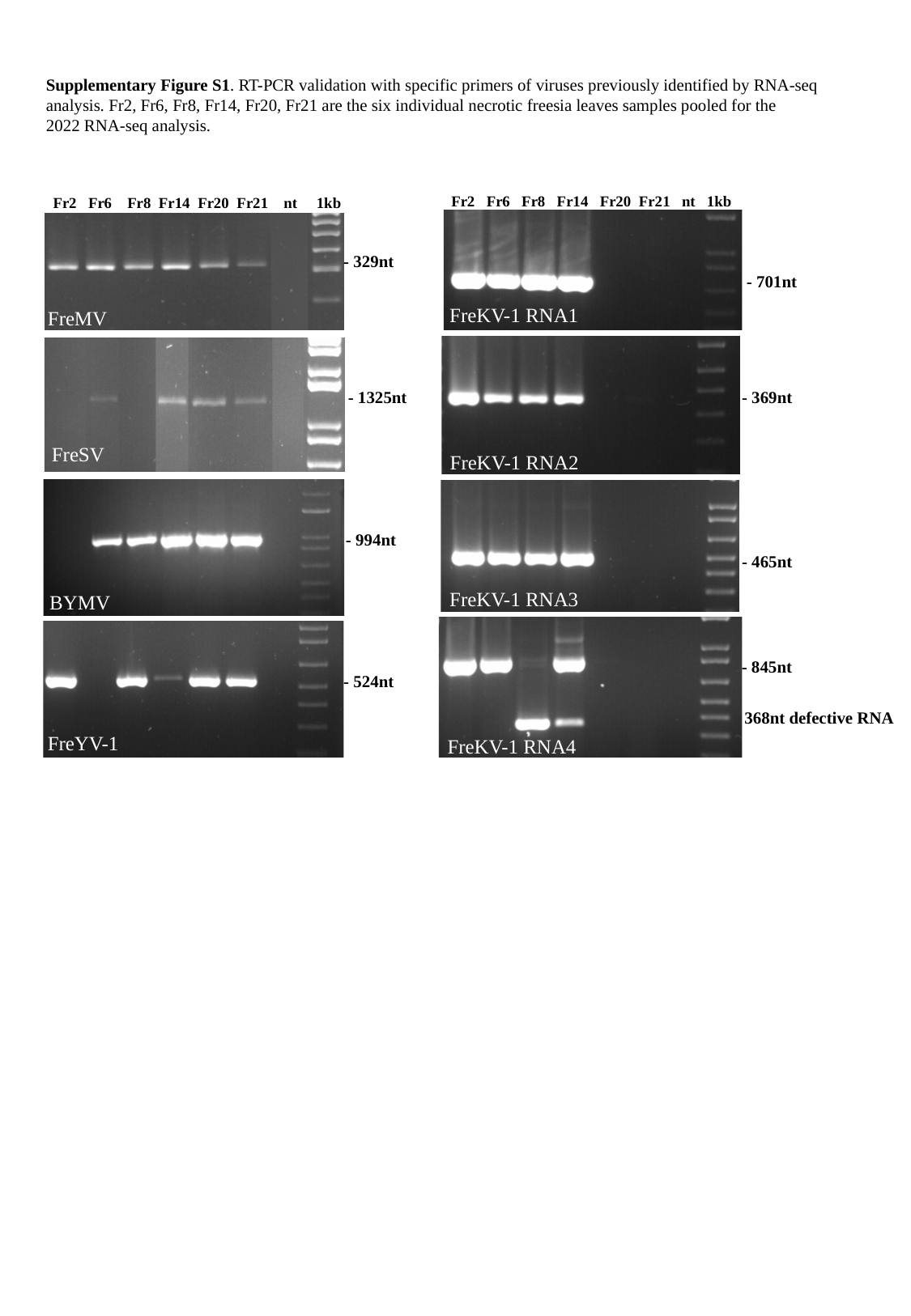

Supplementary Figure S1. RT-PCR validation with specific primers of viruses previously identified by RNA-seq analysis. Fr2, Fr6, Fr8, Fr14, Fr20, Fr21 are the six individual necrotic freesia leaves samples pooled for the 2022 RNA-seq analysis.
 Fr2 Fr6 Fr8 Fr14 Fr20 Fr21 nt 1kb
- 701nt
FreKV-1 RNA1
 Fr2 Fr6 Fr8 Fr14 Fr20 Fr21 nt 1kb
- 329nt
FreMV
- 369nt
FreKV-1 RNA2
- 465nt
FreKV-1 RNA3
- 845nt
368nt defective RNA
FreKV-1 RNA4
- 1325nt
FreSV
- 994nt
BYMV
- 524nt
FreYV-1
