## Supplementary Figure S2 for "The *Freesia refracta* virome analysis sheds new light on the phylogenetic relationships in the *Konkoviridae* and *Yueviridae* families"

### Slide 1
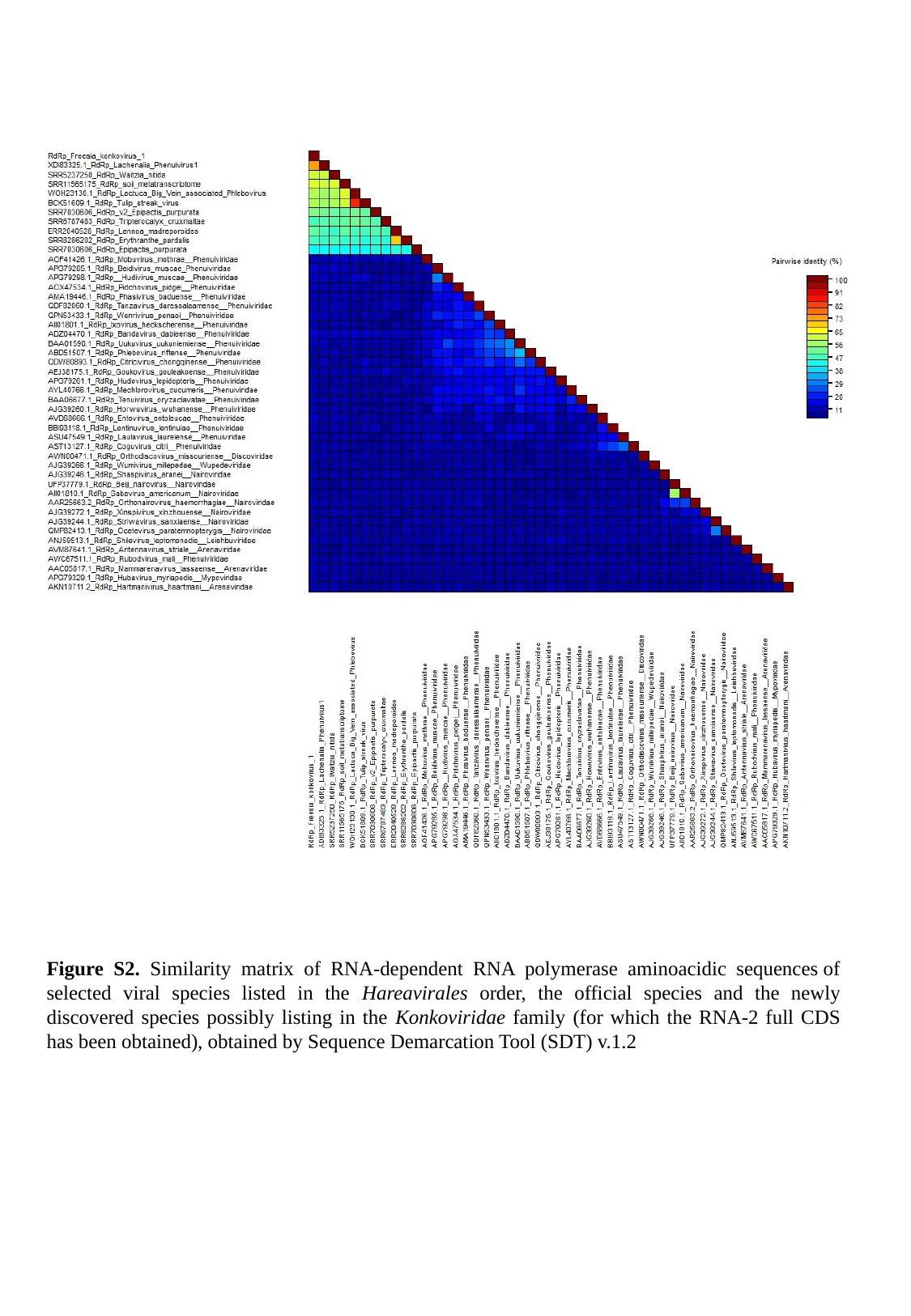

Figure S2. Similarity matrix of RNA-dependent RNA polymerase aminoacidic sequences of selected viral species listed in the Hareavirales order, the official species and the newly discovered species possibly listing in the Konkoviridae family (for which the RNA-2 full CDS has been obtained), obtained by Sequence Demarcation Tool (SDT) v.1.2
