## Supplementary Figure S3 for "The *Freesia refracta* virome analysis sheds new light on the phylogenetic relationships in the *Konkoviridae* and *Yueviridae* families"

### Slide 1
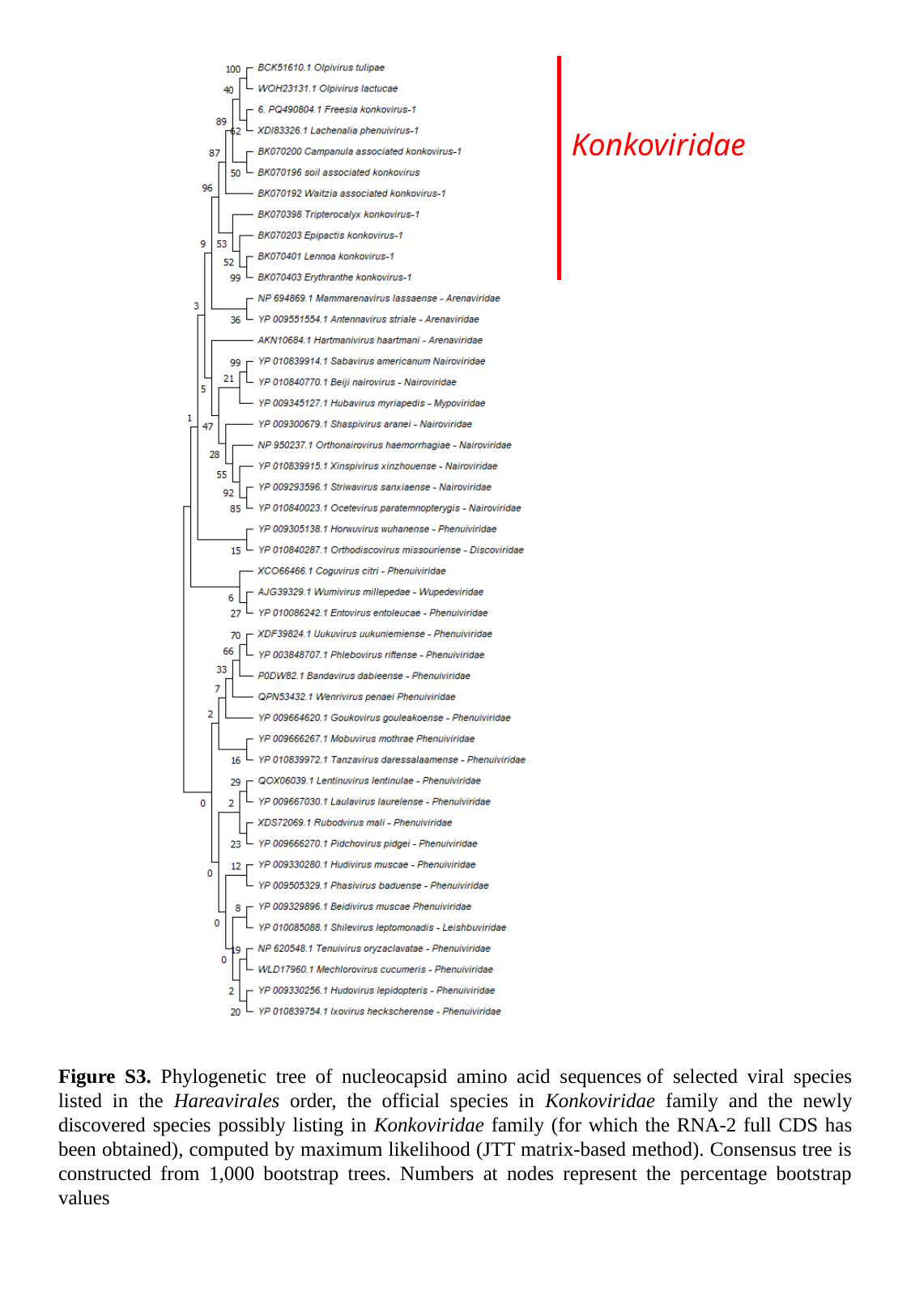

Konkoviridae
Figure S3. Phylogenetic tree of nucleocapsid amino acid sequences of selected viral species listed in the Hareavirales order, the official species in Konkoviridae family and the newly discovered species possibly listing in Konkoviridae family (for which the RNA-2 full CDS has been obtained), computed by maximum likelihood (JTT matrix-based method). Consensus tree is constructed from 1,000 bootstrap trees. Numbers at nodes represent the percentage bootstrap values
