## Supplementary Table S1 for "The *Freesia refracta* virome analysis sheds new light on the phylogenetic relationships in the *Konkoviridae* and *Yueviridae* families"

| **FreKV-1 primers** | **Sequence 5’-3’** | **target** | **Amplified fragment (bp)** | **Source/use** |
| --- | --- | --- | --- | --- |
| TP5-F | GTGGCAGAGTGCTTGCCAAAT | RNA-1 | 701 | Vaira *et al.,* 2018 – diagnosis/validation |
| TP3-R | GGTTCTCCTATCCAAGCACAAGC |  |  |  |
| PhlFRL-412rev | GGAATCTGGCTCTCAAGATG | RNA-1 | cDNA synthesis | 5’RACE-PCR |
| PhlFRL-249rev | TCTCAGGTTTTGTCTGGTGAC | RNA-1 | c.a. 250* | 5’RACE-PCR |
| PhlFR-5853fw | GGCCCACTCCCATTATCATG | RNA-1 | cDNA synthesis | 3’RACE-PCR |
| CNTG271-3race | TAAAAACTTCGCGTCCTTTCAGCA | RNA-1 | c.a. 100* | 3’RACE-PCR |
| NC-F | CCTTCAAAGGCTTCAATCCA | RNA-2 | 369 | diagnosis/validation |
| NC-R | CAGTCCAACGCAATCTCTGA |  |  |  |
| PhlFRS_408rev | AACTGCAACTTGGCCTGAAG | RNA-2 | cDNA synthesis | 5’RACE-PCR |
| PhlFRS_299rev | GAGGCTCCAGAGAAACCAAA | RNA-2 | c.a. 400* | 5’RACE-PCR |
| PhlFRS_549fw | CAGTCCAACGCAATCTCTGA | RNA-2 | cDNA synthesis | 3’RACE-PCR |
| CNTG14-3race | TTGGATTGAAGCCTTTGAAGGAAA | RNA-2 | c.a. 200* | 3’RACE-PCR |
| FreNaV_RNA3_5pfw | ACACAAAGACCGACT**R**C**M**A | RNA-3 | 465 | validation of 5’ end/diagnosis |
| FreNaV_RNA3_470rv | GTGTGTCCAATCGTGCTAGTC |  |  |  |
| FreNaV_RNA4_8fw | GACCGCCCCATCAATTTTCA | RNA-4 | 846 | diagnosis/validation |
| FreNaV_RNA4_853rv | GGAGAGCTCATAACATCATGCC |  |  |  |
| **FreMV primers** |  |  |  |  |
| FreMV-Fw | ACCGCATTTGTTAGCAAGGA | RNA | 329 | diagnosis |
| FreMV-Rv | TCGCCGACACCTTTAAAATC |  |  |  |
| **FreSV primers** |  |  |  |  |
| FOV-CP-Forw | CAGGATCCGATGTCTGGAAAATACTC | RNA3 | 1308 | Rotunno *et al.,* 2024 - diagnosis |
| FOV-CP-Rev | GCGAATTCTTATTAGATAGTGAATCC |  |  |  |
| **FraYV-1 primers** |  |  |  |  |
| ParPol-FW | TTTTACGCGGCTAGTTCACC | RNA-L | 524 | diagnosis/validation |
| ParPol-1-RV | TGAGTTGGTGTACGCTGCTC |  |  |  |

**Supplementary Table S1**. Primers used in this study

*paired with Oligo dT Anchor primer
